# Genome structure and evolution of *Antirrhnum majus* L

**DOI:** 10.1101/443515

**Authors:** Miaomiao Li, Dongfen Zhang, Hui Zhang, Qiang Gao, Bin Ma, Chunhai Chen, Yingfeng Luo, Yinghao Cao, Qun Li, Yu’e Zhang, Han Guo, Junhui Li, Yanzhai Song, Yue Zhang, Lucy Copsey, Annabel Whibley, Yan Li, Ming qi, Jiawei Wang, Yan Chen, Dan Wang, Jinyang Zhao, Guocheng Liu, Bin Wu, Lili Yu, Chunyan Xu, Jiang Li, Shancen Zhao, Yijing Zhang, Songnian Hu, Chengzhi Liang, Ye Yin, Enrico Coen, Yongbiao Xue

## Abstract

Snapdragon (*Antirrhinum majus* L.), a member of Plantaginaceae, is an important model for plant genetics and molecular studies on plant growth and development, transposon biology and self-incompatibility. Here we report a high-quality genome assembly of *A. majus* cultivated JI7 (*A. majus* cv.JI7) of a 510 Mb with 37,714 annotated protein-coding genes. The scaffolds covering 97.12% of the assembled genome were anchored on 8 chromosomes. Comparative and evolutionary analyses revealed that Plantaginaceae and Solanaceae diverged from their most recent ancestor around 62 million years ago (MYA). We also revealed the genetic architectures associated with complex traits such as flower asymmetry and self-incompatibility including a unique TCP duplication around 46-49 MYA and a near complete *ψS*-locus of ca.2 Mb. The genome sequence obtained in this study not only provides the first genome sequenced from Plantaginaceae but also bring the popular plant model system of *Antirrhinum* into a genomic age.

## Introduction

The genus *Antirrhinum* belongs to the Plantaginaceae family and includes about 20 species with the chromosome number of 2n=16^1^. The cultivated *A. majus* was domesticated as an ornamental in garden over two thousand years ago. *Antirrhinum* originated in Europe and are mainly distributed in Europe, Asia, and Africa around the Mediterranean coast. Different species in *Antirrhinum* have obvious differentiation in flower color, fragrance, flower pattern and flowering time; and interspecific hybridization often occurred among them. The genus has evolved two major mating systems, insect pollination (entomophily) and self-incompatibility (SI), to promote out-cross^1,2,3^.

*Antirrhinum* has served as an excellent system in molecular genetics and developmental biology studies in the past three decades because of its active transposable elements for generating rich mutant resources^4^. Several key genes were first cloned in *Antirrhinum* including the founding members of MADS (*DEFICIENS*) and TCP (*CYCLOIDEA*) gene families, a MYB gene *MIXTA* controlling petal epidermis formation, three MYB transcription factors encoded by *ROSEAL*, *ROSEAL2* and *VENOSA* controlling flower color intensity and the *SLF*s (***S***-***L***ocus ***F***-box) controlling self-incompatibility^5-13^. Most of these genes were identified by a representative “cut and paste” transposon-tagging systems in *Antirrhinum* involved in flower development, floral-organ identity and inflorescence architecture ^1,14^.

However, it remains unclear what constitutes their genomic architectures and how they evolve without the genome structure of the regions containing these genes. Here, we report a high-quality genome sequence of *A. majus* of 510 Mb assembled with 37,714 annotated protein-coding genes combining using whole-genome shotgun sequencing of Illumina short reads and single-molecule, real time (SMRT) sequencing long reads from Pacific Biosciences (PacBio) platform. Most of the assembled sequences were anchored to chromosomes to form 8 pseudomolecules using a genetic map. Furthermore, comparative genomic analysis revealed that *Antirrhinum* was derived from related families about 62 MYA and a whole genome duplication event occurred around 46-49 MYA. We also showed a near complete genomic structure of the pseudo (ψ) *S*-locus of *A. majus* of ca. 2 Mb, which consists of 102 genes from *RAD* to *SLF37* genes^15^. The genome sequence provided in this study will accelerate genomic and evolutionary studies on this classical model species.

## Results

### Genome sequencing, assembly and annotation of *A. majus*

We sequenced a highly inbred *Antirrhinum* line (*A. majus* cv. JI7 with eight linkage groups) using the combination Illumina short read and Pacific Biosciences (PacBio) long read sequencing technologies by the genotyping-by-sequencing (GBS) method^16^. The genome size was estimated to be about 520 M based on kmer distribution. We used Canu to correct and assemble PacBio reads into contigs and using SSPACE for scaffolding with Mate-paired short reads. The assembled genome size was 510 Mb with a contig and scaffold N50 sizes of 733 kb and 3,742 kb, respectively (Figure 1, Table 1, Supplementary Figure1 and Supplementary Table 1-2). To anchor *Antirrhinum* genome sequence to chromosomes, we used JoinMap 4.1 to designed marks from re-sequencing 48 recombinant inbred lines (RIL) that derived from *A. majus* crossed to the self-incompatible species *A. charidemi*. Eight linkage groups covered 496.9Mb, representing 97.12% of the assembled *Antirrhinum* genome. The largest chromosome 2 was estimated ca. 75.4 Mb, whereas the smallest chromosome 4 ca. 50.9 Mb and chromosome 8 ca. 57.0 Mb. The average ratio of physical to genetic distance was estimated to be 753.6 kb/cM. The rate of genetic and physical distance showed lower recombination rate at the pericentromeric regions of chr4, chr6, chr7 and chr8, together with the locations of their centromeres. Validations by known genetic markers and FISH (fluorescence *in situ* hybridization) showed that the linkage groups represent a high-quality physical map (Supplementary Figure 2-4 and Supplementary Table 3).

**Figure 1.**
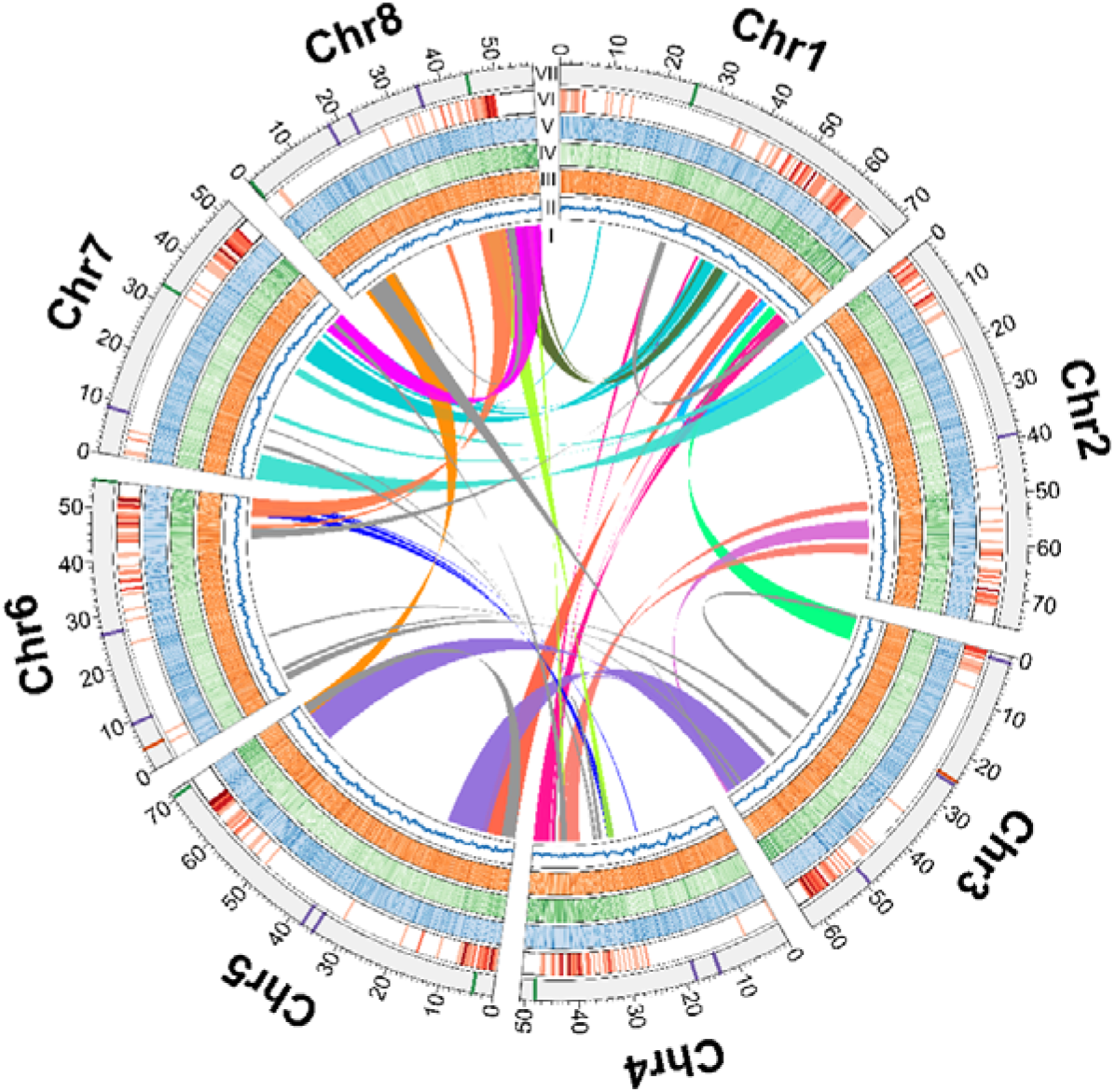
Genomic structure overview of *Antirrhinum majus* JI7. (I) Duplications of genomic paralogous sequences; (II) GC content. In II, III, IV, and V, deep colors show high density genes or repeat sequence regions; (III) Simple sequence repeats; (IV) Gene density; (V) Retroelement density; (VI) Recombination rate. Deep color shows high recombination rates.); (VII) Eight Antirrhinum chromosomes with physical distances including low copy number repetitive elements: telomere repeat TTTAGGG (green), 5S rDNA (orange), pericentromeric repeats CentA1 and CentA2 (purple).

**Table 1.** Statistics for the Antirrhinum genome and gene annotation.

|  |  |
| --- | --- |
| Estimate of genome size | 520 Mb |
| GC content | 35.50% |
| N50 length (contig) | 0.73 Mb |
| Longest contig | 3.74 Mb |
| Total size of assembled contigs | 510.00 Mb |
| N50 length (scaffold) | 2.62 Mb |
| Longest scaffold | 9.90 Mb |
| Total size of assembled scaffolds | 511.70 Mb |
| Number of genes | 37714 |
| Average gene length | 3166 bp |
| Gene density | 73.95 Mb<sup>-1</sup> |
| Transcripts number | 52780 |
| Average CDS length | 1036 bp |
| Average protein length | 344 aa |
| Average exon length | 245 bp |
| Average intron length | 314 bp |
| Tandem repeat | 13.03 Mb |

To validate genome quality and integrity, the completeness of the assembled euchromatic portion was examined by comparing 25,651 public ESTs of *Antirrhinum* from NCBI (http://www.ncbi.nlm.nih.gov/nucest/?term=EST%20Antirrhinum) and 96.59% EST sequences were mapped on the assembled genome. Next, we did the alignments between three Bacterial Artificial Chromosome (BAC) sequences and the assembled *Antirrhinum* genome, indicating an average 98.3% of euchromatin coverage. We also used BUSCO analysis to compare the quality of the assembled *Antirrhinum* genome sequence to that of other published plant genomes and showed a high quality of the assembled sequence data (Supplementary Figure 5-7).

We predicted a total of 37,714 protein-coding genes with an average transcript length of 3,166 bp by using *Antirrhinum* EST sequences and RNA-seq data from six major tissues: leaf, root, stem, stamen, pistil and pollen (http://bioinfo.sibs.ac.cn/Am/, Supplementary Excel 1). Approximately 89.11% of all genes could be functionally annotated. The average gene density in *Antirrhinum* is one gene per 15.5 kb, which is about three times lower than Arabidopsis (one gene/4.5 kb) and slightly higher than tomato (one gene/25.7 kb). Genes are also distributed unevenly, being more abundant in the ends of chromosomal arms. We also identified 981 transfer RNA, 800 microRNA, 10 ribosomal RNA and 622 small nuclear RNA families. A total of 268.3 Mb (43.8%) sequences were annotated as repeats including a wealth of class I (Retrotransposon: 182.8 Mb), class II (DNA transposon: 41.1 Mb) elements, accounting for 52.6% of the assembled genome (Supplementary Table 4-7).

To detect the intragenomic organization in *A. majus*, self-alignment analysis was employed to reveal the duplicated and triplication regions between and within chromosomes. A whole genome triplication occurred between chromosome 1, 7 and 8, between chromosome 4, 6 and 8. Paralogous relationships among eight *Antirrhinum* chromosomes revealed 45 major duplications and 2 triplications by dot plot analyses. Collectively, forty-seven major intra-syntenic blocks spanning 1,841 pairs of paralogous genes were identified on the eight chromosomes (Figure 1, Supplementary Excel 2).

Taken together, the *Antirrhinum* genome generated in this study represents a high-quality chrome-scale genome map.

### Comparative genomics and evolution of *A. majus*

To compare *Antirrhinum* genome with other plant genomes, we first examined the synteny of *Antirrhinum* chromosomes. The results showed that homologous genes in *Antirrhinum* chromosomes 1, 4, 5, 6, 7 and 8 are collinear to tomato chromosome 6/7/11, 3/8, 1/4/5, 2/3, 12 and 9, respectively, indicating that these chromosomes shared similar evolutionary histories (Figure 2a). Base on the collinearity of gene pairs, we calculated the density distribution of synonymous substitution rate per gene (*Ks*) between collinear paralogous genes and inferred the time of whole-genome duplication (WGD) events in *Antirrhinum*. A peak at around 0.57-0.60 showed a recent WGD in *Antirrhinum* occurred around 46-49 Mya corresponding to a β event ^17^ (Figure 2b). We then compared the complexity of gene families between *Antirrhinum* and other species and showed that 9,503 gene families are shared by *Antirrhinum*, *Arabidopsis*, rice, and tomato. 6,677 genes families were possibly contracted in *Antirrhinum* while the other 3,778 gene families expanded including the F-box protein family being expanded significantly (Figure 2c).

**Figure 2.**
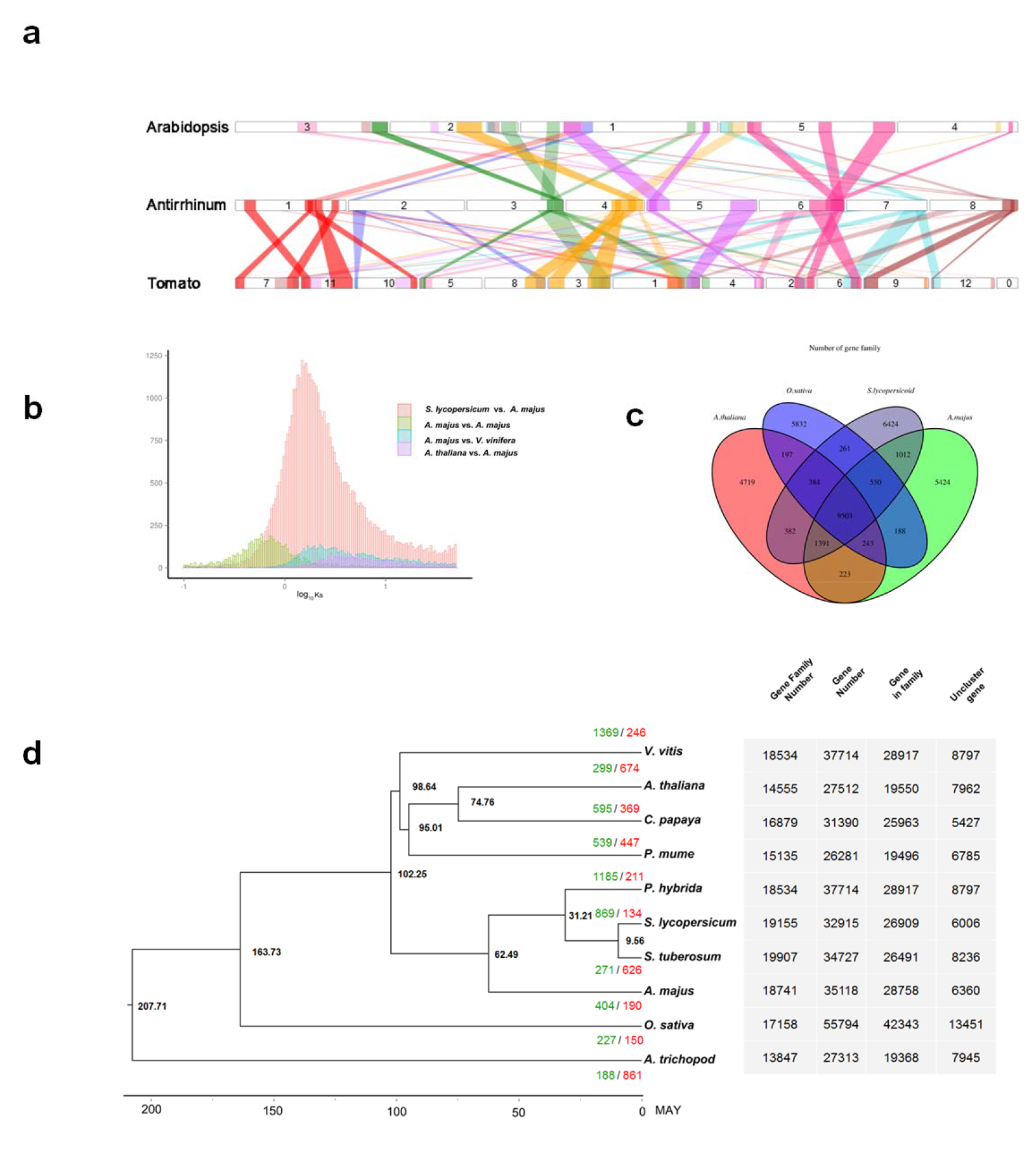
Genome evolution of *A. majus*. **a** Synteny blocks among chromosomes of *A. majus*, *S. lycopersicum* and *A. thaliana.* The numbers represent individual chromosomes. The selected syntenic gene number more than fifty. **b** Density distributions of Ks for paralogous genes. **c** A Venn diagram of shared orthologues among four species. Each number represents a gene family number. Venn diagram of annotated gene families showing shared orthologous groups among the genomes of *A. majus*, *S. lycopersicum*, *A. thaliana*, and *O. sativa*. **d** Phylogenetic tree of 10 angiosperm species including their divergence time based on orthologues of single-gene families. The number in each node indicates the number of gene families. *A. trichopoda* used as an outgroup. Bootstrap values for each node are above 100%.

To examine the *Antirrhinum* genome evolution, we performed all-against-all comparison analyses for the evolution of gene families and constructed a phylogenetic tree of nine angiosperm species (*A. majus*, *A. thaliana*, *A. trichopoda*, *C. papaya*, *O. sativa*, *P. hybrida*, *P. mume*, *S.lycopersicum*, *S. tuberosum* and *V. vitis*) based on the shared 2114 single-copy genes of *Antirrhinum*. The divergence times of *C. papaya***-***A. thaliana* (55.1~90.6 million years ago) and dicot-monocot (123.9~228.5 million years ago) were used for calibration derived from the published data (Figure 2d). Taken together, these results showed that *Antirrhinum* lineage was split from potato and tomato lineages around 97 MYA with its own recent WGD event occurred about 46-49 MYA.

### Evolution of floral asymmetry and TCP family

*A. majus* has served as the genetic model of floral symmetry. Recent studies have revealed the floral asymmetry in *A. majus* is largely controlled by two TCP family TFs (CYC and DICH) ^7,8,18^. To explore their evolution, we compared the composition and number of TCP family in several sequenced angiosperms with floral symmetry information. Both eudicot and monocot share a CYC/DICH embedded CYC/TB1 clade, while radial flower basal angiosperm *A. trichopoda* lacks any members of class II CYC/TB1 clade, suggesting that class I and class II CIN clade were more ancient than class II CYC/TB1 clade, and the initial role for CYC/TB1 clade was not likely involved in floral symmetry control (Figure 3 and Supplementary Excel 3).

**Figure 3.**
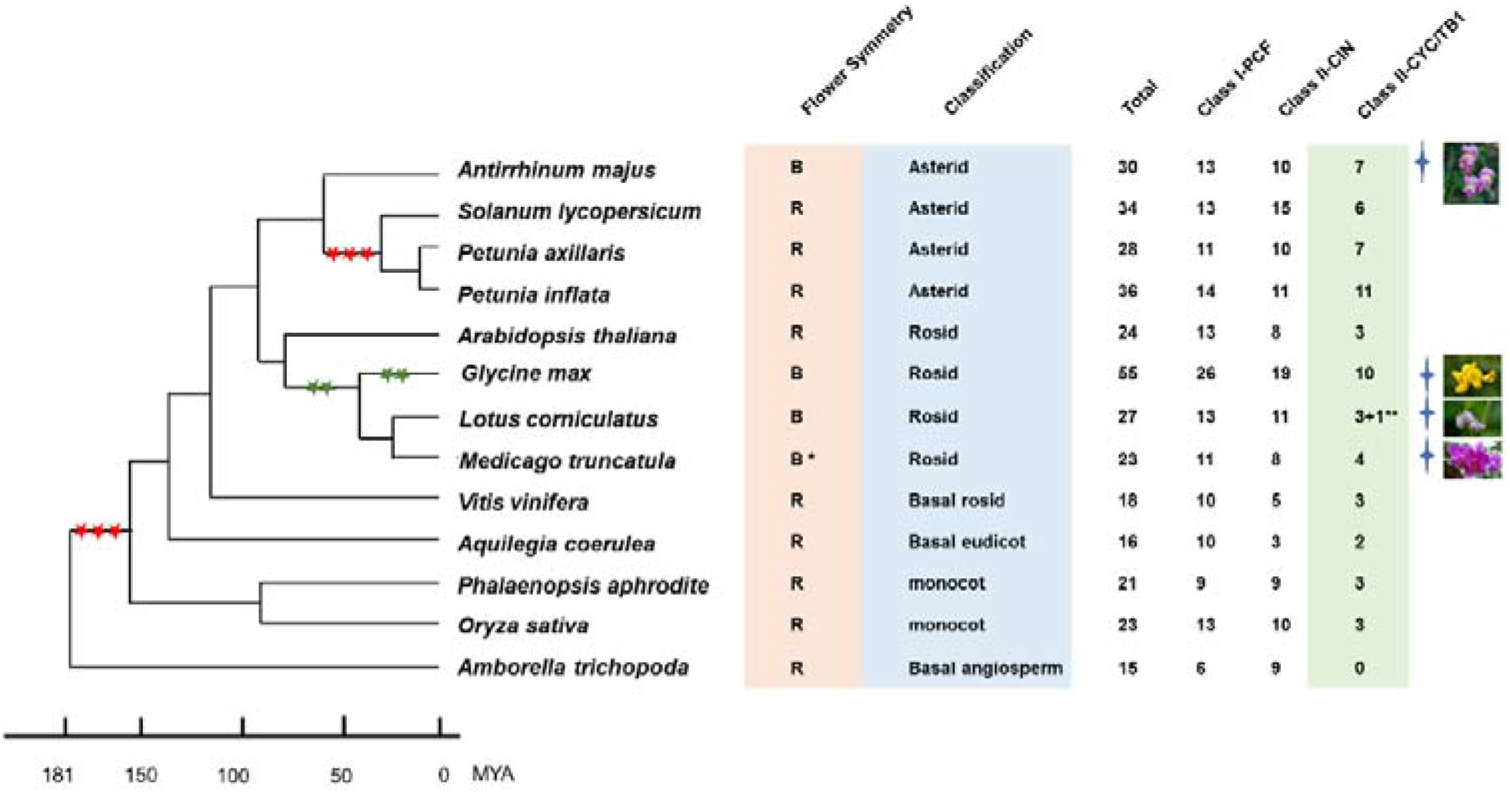
Evolution of flower symmetry and TCP gene family. Left shows a phylogenetic tree of the flowering plants derived from their divergence time based on orthologues of single-gene families. Three red stars show the whole genome triplication and two green stars the duplication events. “B” represents bilateral flower symmetry and “R” radial flower symmetry; Asterid, Rosid, Basal rosid, Basal eudicot, Monocot and Basal angiosperm represent their clades, respectively. Total numbers of TCP family genes, Class I-PCF, Class II-CIN and Class II-CYC/TB1 are shown from left to right. * indicates the sequenced genome of species of *Medicago truncatula* with flower radial symmetry, but flowers of most *Medicago* species are of bilateral symmetry. ** indicates *Lotus comiculatus* in which three TCP genes were identified but a functional TCP gene was not detected in its genome. Four-pointed stars denote bilateral symmetry flowers with their photos from PPBC (http://www.plantphoto.cn).

We identified a total of 30 putative functional TCP family genes, including 13 class I genes, 10 class II CIN clade genes and 7 class II CYC/TB1 clade genes. Syntenic block and *Ks* analyses of the orthologous gene pairs revealed that the recent whole genome duplication and tandem duplication contributed to the expansion of TCP family members, especially for the *CYC*/*DICH* embedded CYC/TB1 clade (Figure 3 and Supplementary Table 8). Previous studies indicated *CYC* and *DICH* have overlapping expression patterns in floral meristems in *A. majus* and the fully radial flower only appeared in *CYC*/*DICH* double mutants, suggesting the genetic robustness of zygomorphic flower control. To estimate the time of *CYC* and *DICH* duplication event, we performed *Ks* analysis of the syntenic block embedding *CYC*/*DICH* with 79 homologous gene pairs (Supplementary Table 8). Interestingly, the results indicated this syntenic block was retained from a whole genome duplication event taken place during the time span of the zygomorphic flower emergences (about 60 million years ago). Furthermore, downstream *CYC/DICH*, two genes (*DIV* and *DRIF*) in *RAD*/*DIV* module have homologous copies with similar Ks as *CYC*/*DICH*, while *DRIF* is located at a WGD derived syntenic block^19,20,21^. Those analyses showed that the master regulators of zygomorphic flower were retained from the recent whole genome duplication, revealing the evolutionary base for the zygomorphic flower maintenance in *A. majus* and perhaps other close relatives.

To further explore TCP family evolution among other zygomorphic flower species, we extended the syntenic block and *Ks* analyses to *M. truncatula*, belonging to legume^22^. The results showed that *M. truncatula* independently undergone a recent whole genome duplication event around the similar time as *A. majus* without similar duplicated *CYC*/*DICH* copies, suggesting that *A. majus* harbored a unique genetic mechanism for zygomorphic flower (Supplementary Figure 8). Though we could not couple the genetic events with the appearance of zygomorphic flower, our TCP family analysis on representative angiosperms indicated that the genetic origins of zygomorphic flower were ancient and occurred around the late period of Cretaceous about 60 MYA (Figure 3).

Previous phylogenetic analysis suggested that zygomorphic flower independently evolved from actinomorphic ancestors for more than 25 times^23^. Our findings indicated that gene duplications are involved in a genetic robustness of zygomorphic flower evolution by maintaining copies of master regulators.

### Structure of the *ψS*-locus in *A. majus* and its gene collinearity in SI species

Previously, we found the *Antirrhinum S*-locus is located in a heterochromatin region on the short arm of chromosome 8 through cytological investigation^24^. To reveal a complete pseudo (*ψ*)*S*-locus in self-compatible *A. majus*, we scan conserved regions (FBA/FBK domain) of *SLF* gene family in the assembled *A. majus* genome, we identified the *S* genes located in the short arm of chromosome 8 and examined their expressions inside the *ψS-*locus region. The locus consists of 37 *SLF* genes (*SLF1*-*SLF37*) covering 874 kb composed of three scaffolds *Sc29*, *Sc276* and *Sc184* (Figure 4a). Six pseudogenes with FBA domains were inferred to be loss-of-function in evolution. No S-RNase was found in and near the region, suggesting it might be lost in self-compatible *A. majus*. *RAD* gene locates in the upstream of *SLF1* gene about 1Mb, consistent with previous studies showing its linkage with the *S*-locus^24^; and the region contains 102 genes from *RAD* to *SLF37*. Most *SLF* genes are expressed in pollen or anther indicating they could be related to pollination and fertilization. Taken together, these results showed that we obtained a near complete ψ*S*-locus with the largest numbers of *S* genes annotated in a plant genome (Figure 4b and Supplementary Excel 4-6).

**Figure 4.**
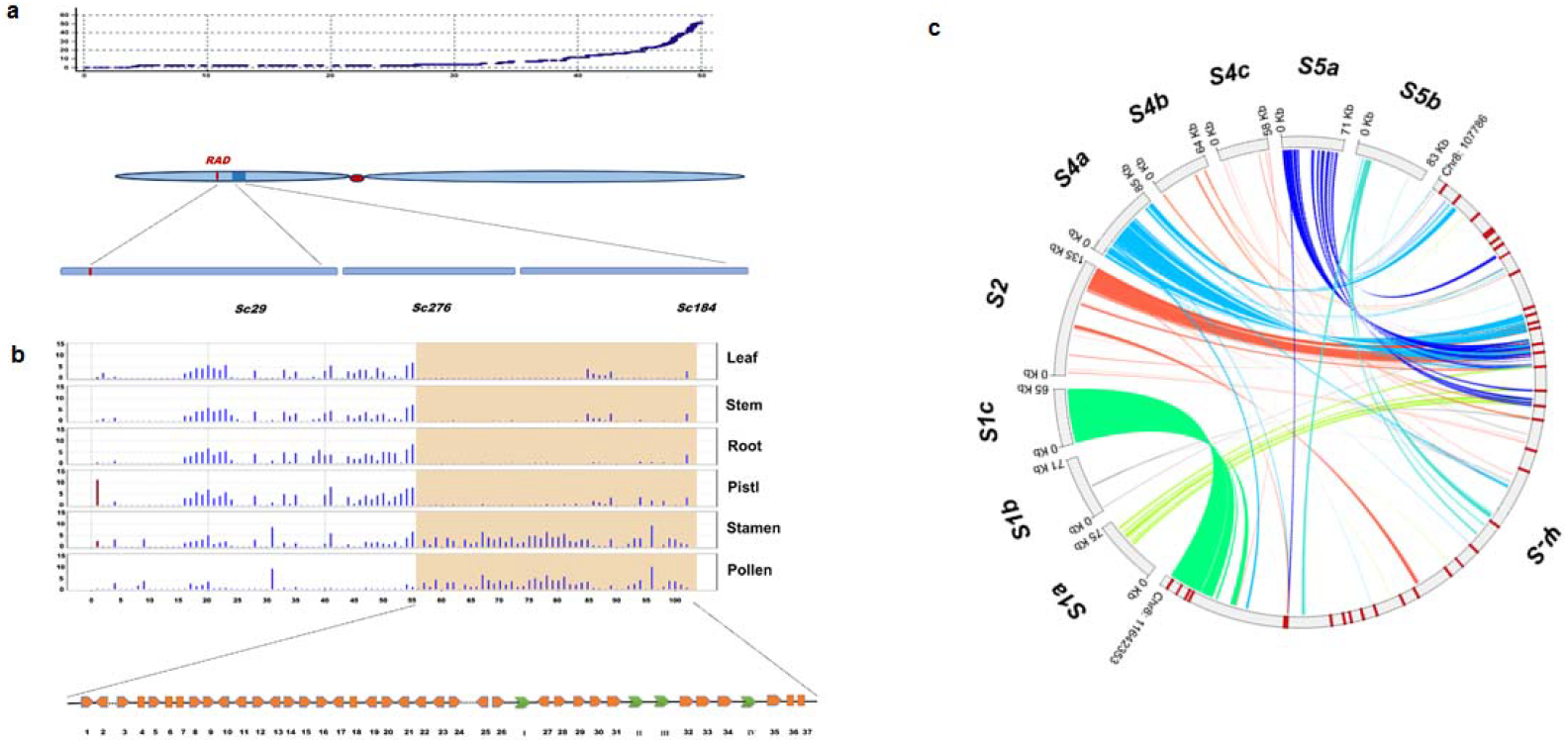
Genomic features of the *ψS*-locus of *A. majus* and its synteny with the *S*-locus regions of *A. hispanicum*. **a** Chromosomal locations of three scaffolds covering the *ψS*-locus region of *A. majus*. A genetic recombination map of chromosome 8 is shown on the top panel. The x-axis shows its physical distance and the y-axis its genetic distance. A schematic representation of chromosome 8 is shown in the middle panel with a red dot indicating its centromere. The *ψS*-locus is depicted as a blue box on its short arm. A vertical red line in chromosome indicates *RAD* gene. The low panel shows three scaffolds of *Sc29*, *Sc276*, and *Sc184* covering the *ψS*-locus region. **b** Transcriptional profiles of the *ψS*-locus and its flanking regions of *A. majus*. The light yellow shadow denotes the predicted *ψS*-locus region (*SLF1*-*SLF37*). This region between *RAD* and *SLF37* contains a total number of 102 annotated genes. The bottom part is a schematic representation of the *SLF* genes. Orange squares indicate the *ψSLF* genes and green arrows the other annotated genes (I: a putative MYB family transcription factor; II and III, putative RNA-binding proteins; and IV, a putative phosphate-dependent transferase) **c** The synteny of the *S*-locus regions between *A. majus* and *S_1_*, *S_2_*, *S_4_* and *S_5_* haplotypes of *A. hispanicum*. Different colors indicate syntenic and inversion regions between the *ψS*-locus and *S_1_* (*S_1a_*, *S_1b_* and *S_1c_*), *S_2_*, *S_4_* (*S_4a_*, *S_4b_* and *S_4c_*) or *S_5_*(*S_5a_* and *S_5b_*) haplotypes of *A. hispanicum*.

To explore the *S*-locus structures between SC and SI species, we compared the *ψS*-locus sequence with nine assembled TAC (Transformation-competent artificial chromosome) sequences from four *S* haplotypes of self-incompatible *A. hispancum* and uncovered high gene collinearity between *AhSLF12* and *AhSLF13* except for *AhSLF32* located in *S_2_*, *S_4_* and *S_5_* haplotypes of *A. hispanicum* (Figure 4c). An intra-chromosome inversion around the *S*-locus occurred in the *S_2_* haplotype of *A. hispanicum* as described previously. However, no apparent large collinearity was found between *A. majus* and TAC sequences containing the *S-RNase* gene, supporting that the region containing *S-RNase* was deleted in the *ψS*-locus. We found one orthologous pseudo-gene in *A. majus*, *AmSLF18* on chromosome 8, which has a complete corresponding coding sequence region and expressed in the *S_4_* haplotype, suggesing a recent duplication event (Figure 4c, Supplementary Excel 7).

To examine the evolution of *SLF and S-RNase* genes, the *Ka* and *Ks* rate of the twelve collinear *SLF* gene pairs showed that the values of *SLFs* are lower than that of *S-RNase* in *Antirrhinum*, and the allelic *SLF* genes showed a Ka/Ks=0.41, consistent with a negative frequency-dependent selection detected previously. Only *SLF14* appears to be a positively selected gene (Ka/Ks>1) (Supplementary Excel 8). The average divergence time of these *SLF* orthologous genes was estimated to be 4 million years ago consistent with the early *Antirrhinum* species divergence of less than 5.3 million years. The average divergence time of these *S-RNases* is estimated to be around 90 MYA, consisting with the species divergence between *Antirrhinum* and Solanaceae species (Supplementary Excel 9).

Taken together, our results showed that the *Antirrhinum S*-locus is shaped by allelic synteny, purifying selection, gene conversion, transposition and pseudogenization.

## Discussion

In this study, we have successfully obtained a chromosome-scale fine genome of *A. majus*, representing the first sequenced genome in Plantaginaceae. This genome provides a useful resource for evolutionary and molecular studies on key genes controlling complex traits in the classical eudicot model species. The comparison of 404 gene families’ expansions and 191 contractions in *A. majus* genome with those of nine other fully sequenced diverse plant species showed several specific evolutionary paths of important gene families, such as those involved in metabolism and signaling pathways. Furthermore, the absence of a recent WGT in *Antirrhinum* genome makes it a key diploid genome that promises to provide important insights into plant genome evolution. Recently, small RNAs in natural hybrid zone of *A. majus* using this genome sequence as a reference have revealed an inverted duplication acting as steep clines by nature selection in evolution of flower color pattern^25^.

Complex traits, such as zygomorphic type flower in *Antirrhinum*, often couple with different gene duplication events. The zygomorphic structure in *Antirrhinum* occurred in late period of Cretaceous, consistent with its role in facilitating insect-mediated pollination. We demonstrated that the morphology of floral organ was associated with several gene duplication events and revealed the divergence of *A. majus* from other angiosperms. Although *Antirrhinum* did not experience a whole genome triplication, the duplicated region containing TCP genes on chromosomes 6 and 8 are closely associated with zygomorphic flower type, showing its unique evolutionary manner because no similar mechanism had been found in closely related species. For examples, one separated whole genome duplication led to partial zygomorphic type species of Glycine; and a MADS box family gene change resulting in symmetrical flowers in Orchidaceae^26,27^. In general, the ancient species are mostly radial symmetry and the newly derived species with symmetrical flowers. Studies on *Antirrhinum* floral mutants supported that these two copies play a role in controlling symmetrical flower, which makes the petals polar, and the flower structure tends to be upright and is beneficial to the use of light and attracting pollinators. This mechanism could be further verified by genetic analysis in a relative species.

SI also is a complex trait controlled by the *S*-locus^2,3^. The fine genomic structure of the *ψS*-locus from *A. majus* showed that the large number of pollen *SLFs* could be a result of recombination suppression, gene duplication, purifying selection and frequency-dependent selection^28^. These evolutionary processes could be the intrinsic mechanisms to maintain the low allelic diversity of orthologous *SLFs* because extensive divergence would lead to self-inactivation of S-RNase resulting in loss of SI. The deletion of S-RNase in cultivated *A. majus* could be directly associated with the loss of SI, resulting in an irreversible evolutionary process, because loss of self-incompatibility of SI species often results in compatible ones and SC species are difficult or almost impossible to reverse back to self-incompatible species (Doll’s Law) ^29^. The high microcollinearity of the *S*-locus between SI and SC of *Antirrhinum* indicated that the deletion of *S-RNase* in SC species was a recent event. Some mutated *SLF* genes in different haplotypes were also evolved recently. Loss of *S-RNase* at the *S*-locus would have little consequence for the pollen phenotype while duplications would lead to expression of two types of *SLFs* which could inactivate a broader range of incoming S-RNases. Further genome analysis of additional SI species of *Antirrhinum* should provide an insight into its evolutionary status.

Intriguingly, the physical size of the *S*-locus in *S. lycopersicum* (17 Mb, containing 17 *SLF* genes) is much larger than that in *A. majus* (2 Mb, containing 37 *SLF* genes) ^30^. This appears to indicate that increasing the gene numbers through in-locus unequal crossovers and repetitive element enrichments by a hitchhike effect could result in its large physical size and low gene density of the *S*-locus in tomato. Rich retrotransposons and unique small regulatory RNAs associated with the *S*-loci of *Solanum* and *Antirrhinum*, respectively, appear to indicate the differential epigenetic modifications could contribute to the locus density puzzle between these species.

In conclusion, the high-quality genome sequence obtained in this work could be used as a reference genome for Plantaginaceae and will be helpful for genetic, genomic and evolutionary studies in both *Antirrhinum* and other flowering plants.

## URLs

Genome assembly data has been deposited at NCBI BioProject ID under accession codes PRJNA227267 and Biosample ID under accession codes SAMN02991092. We built the website of *Antirrhinum* genome http://bioinfo.sibs.ac.cn/Am.

## Methods

Methods and any associated references are available in the online version of the paper.

## Acknowledgements

This work was supported by the Ministry of Science and Technology of China (2013CB945102) and the National Natural Science Foundation of China (31401045 and 31221063). We thank Zhixi Tian for help of CACTA transposon analyses.

## Author contributions

Y.X., H.Z., D.Z. and M.L. designed the experiments; M.L., D.Z. and Y.X. wrote the manuscript. Q.G, B.M., C.C., Y.L., Q.L., Y.Z., H.G., J.L., Y.Z., Y.S., L.C., A.W., Y.C., Y.L., M.Q., J.W., Y.C., D.W., J.Z., G.L., B.W., L.Y., C.X., J.L., S.Z., Y.Z, S.H., C.L., Y.Y., E.C. and Y.X. analyzed the data and performed the experiments.

